# Improvement of D-lactic acid production from methanol by metabolically engineered *Komagataella phaffii* via ultra-violet mutagenesis

**DOI:** 10.1101/2024.12.09.627664

**Authors:** Yoshifumi Inoue, Kaito Nakamura, Ryosuke Yamada, Takuya Matsumoto, Hiroyasu Ogino

## Abstract

Methanol has attracted attention as an alternative carbon source to petroleum. *Komagataella phaffii*, a methanol-assimilating yeast, is a useful host for the chemical production from methanol. A previous study successfully constructed a metabolically engineered *K. phaffii* GS115/S8/Z3 strain capable of producing D-lactic acid from methanol. In this study, we aimed to develop a strain with improved D-lactic acid production by applying ultra-violet mutagenesis to the D-lactic acid-producing strain, GS115/S8/Z3. The resulting mutant strain DLac_Mut2_221 produced 5.38 g/L of D-lactic acid from methanol, a 1.52-fold increase compared to the parent strain GS115/S8/Z3. Transcriptome analysis revealed 167 differentially expressed genes in DLac_Mut2_221, comprising 104 upregulated and 63 downregulated genes. These results suggest that the improvement in D-lactic acid production from methanol in the mutant strain involves three important mechanisms: (1) avoiding excessive formaldehyde accumulation, (2) activating the glyoxylate pathway, and (3) reducing the expression of *O*-glycosylated proteins required for cell wall stability. Metabolic engineering strategies for *K. phaffii* based on the knowledge gained from this study will contribute to improving the productivity of various useful chemicals from methanol.

## 1. Introduction

Methanol, a C1 compound, continues to undergo advances in manufacturing technology (Deka et al., 2022). Additionally, the synthesis of methanol from CO_2_ and waste biomass is actively progressing, and the methanol market is expected to continue to grow (Schrader et al., 2009; Kelso et al., 2022). Furthermore, compared to other C1 compounds, such as CO_2_ and methane, methanol has the advantage of being a liquid at room temperature, facilitating easier transportation (Wang et al., 2020). These characteristics position methanol as a promising alternative carbon resource to petroleum. Currently, the production of useful compounds using methanol relies mainly on chemical synthesis in which methanol is first converted into ethylene or propylene, followed by the synthesis of various useful compounds (Tabibian and Sharifzadeh, 2023). These chemical processes require high-temperature and high-pressure conditions and have low reaction specificity, requiring huge energy and resource costs to produce the products (Burg et al., 2016).

In contrast to chemical processes, bioprocesses have several advantages: they can operate under mild conditions, exhibit high reaction specificity, allow the synthesis of structurally complex compounds and polymers, and allow selective production of chiral compounds (Süntar et al., 2021). Owing to these benefits, research on the production of valuable compounds from methanol using methylotrophic microorganisms has increased (Zhang et al., 2018; Zeng, 2019; Chen and Lan, 2020). Lactic acid is particularly advantageous for bioprocess production. As chiral compounds, both L- and D-forms are utilized in various fields, including the food, cosmetics, pharmaceutical, and chemical industries (Santoro et al., 2016; Jia et al., 2018; Esteban and Ladero, 2018; Beitel et al., 2020; Alexandri et al., 2022). Notably, polylactic acid as a bioplastic has garnered significant attention, with studies reporting improved polymer thermal resistance when equal amounts of poly L-lactic acid and poly D-lactic acid are combined to form a stereocomplex structure (Tsuji, 2005). However, only a limited number of lactic acid bacteria can selectively produce D-lactic acid over L-lactic acid, and there is a need to establish a highly efficient method for producing D-lactic acid (Klotz et al., 2016).

*Komagataella phaffii* (formerly *Pichia pastoris*) is a safe microorganism and can be genetically modified. It can metabolize methanol as its sole carbon source and has been used as a host for valuable protein and compound production (Ahmad et al., 2014; Wu et al., 2023; Zhang et al., 2023). Lactic acid can be produced from methanol using methylotrophic yeasts (Yamada et al., 2019; Wefelmeier et al., 2023; Wu et al., 2025). In our previous study, we constructed *K. phaffii* GS115/S8/Z3, in which multiple copies of the *DLDH* gene derived from *Leuconostoc mesenteroides* were introduced into the rDNA locus, and the strain produced 3.48 g/L D-lactic acid from methanol (Yamada et al., 2019). However, these levels are significantly lower than those produced from glucose (Mitsui et al., 2020; Pangestu et al., 2022). Methanol taken up by the cell is first converted to formaldehyde by alcohol oxidase (*AOX*), which is then converted to pyruvate via the xylulose-5-phosphate pathway during assimilation metabolism (Jordà et al., 2012; Unrean, 2013). Lactic acid is produced by the conversion of pyruvate by lactate dehydrogenase (*LDH*). During the dissimilation of methanol in methylotrophic yeasts, formaldehyde is converted to formic acid, which is excreted as CO_2_. To promote the production of methanol-based compounds, suppressing this dissimilation pathway (Guo et al., 2021), avoiding the accumulation of formaldehyde (Zhang et al., 2003; Pedro et al., 2015), and enhancing the metabolic pathways of target compounds have been suggested (Wu et al., 2023). However, methanol metabolism in *K. phaffii* is complex and tightly regulated, and metabolic engineering techniques to efficiently convert methanol into useful compounds are still under development.

Ultra-violet (UV) mutagenesis randomly alters the DNA base pairs in the genome of an organism. UV mutagenesis is a simple method of irradiating organisms with UV radiation, which can damage genomic DNA and introduce mutations during repair. Owing to its simplicity and usefulness, numerous studies on various organisms have used UV irradiation to generate strains with desirable phenotypes (Moshinsky and Wogan, 1997; Guo et al., 2019; Karitani et al., 2024). In *K. phaffii*, UV irradiation improves the production of rFIP-glu protein (Wu et al., 2021). These findings suggest that by introducing UV mutations into *K. phaffii*, it may be possible to obtain mutant strains with enhanced methanol utilization pathways that are intricately regulated.

In this study, we aimed to develop a *K. phaffii* strain capable of producing D-lactic acid from methanol with high efficiency. UV irradiation was performed on the metabolically engineered D-lactic acid-producing strain *K. phaffii* GS115/S8/Z3 (Yamada et al., 2019) to obtain mutant strains with improved D-lactic acid production. Additionally, transcriptome analysis was conducted on the mutant strain to investigate genes related to D-lactic acid production from methanol in *K. phaffii*.

## 2. Materials and Methods

### 2.1 Strains and media

The metabolically engineered D-lactic acid-producing strain, *K. phaffii* GS115/S8/Z3 (Yamada et al., 2019) was used in this study. Yeast/peptone/dextrose (YPD) medium containing 10 g/L yeast extract (Formedium, Norfolk, UK), 20 g/L peptone (Formedium), and 20 g/L glucose (Nacalai Tesque, Kyoto, Japan) and yeast/peptone/methanol (YPM) medium containing 10 g/L yeast extract, 20 g/L peptone, and 30 g/L methanol (Nacalai Tesque) were used for cultivation. If necessary, 20 g/L agar (Nacalai Tesque) and 0.1 g/L zeocin antibiotic (InvivoGen, California, USA) were added.

### 2.2 Yeast cultivation

Microplate culture was performed in 1.0 mL of YPM medium using a 2-mL 96-well deep-well plate equipped with a gas-permeable seal (EXCEL Scientific, California, USA) and a rotary plate shaker (Taitec, Nagoya, Japan) set at 30°C and 1,200 rpm. Cultivation was initiated by culturing the cells in a well containing 1.0 mL of YPD medium at 30°C and 1,200 rpm for 24 h, harvesting and washing the cells, and suspending them in 1.0 mL of fresh YPM medium.

Flask cultures were performed using a rotary shaker (Taitec) operated at 30°C, 200 rpm, with 250 mL flasks containing 50 mL of YPM medium equipped with a gas permeable seal (EXCEL scientific). Cultures were started by inoculation (initial OD_600_ of 10.0) of pre-cultures grown in 250 mL flasks containing YPD medium for 24 h at 200 rpm and 30°C.

### 2.3 Mutagenesis into *K. phaffii*

Yeast cells were cultured in the YPD medium for 24 h. After centrifugation, the cell pellet was resuspended in sterile water to an OD_600_ of 0.5 in a total volume of 10 mL. The suspension was transferred to a petri dish and irradiated with UV light at 18 J/m^2^ for 77 min while stirring. This duration corresponds to the time required to reduce the cell viability to ∼5% (data not shown). After UV exposure, the suspension was collected and plated onto YPD plates containing 0.1 g/L zeocin. Colonies formed after 72 h of incubation at 30°C were isolated and maintained on fresh YPD agar plates containing 0.1 g/L zeocin.

### 2.4 Analysis of growth and metabolites

The OD _600_ of each culture was determined using a spectrophotometer (Shimadzu, Kyoto, Japan).

Methanol and D-lactic acid concentrations were measured using a microplate reader (Thermo Fisher Scientific, Waltham, MA, US) following a previously reported method (Inoue et al., 2024).

### 2.5 Transcriptome analysis

Total RNA was extracted from the cells after 72 h of culture and transcriptome analysis was performed. The cells were collected from the culture broth by centrifugation and disrupted using zirconia beads. Total RNA was isolated from the homogenate using a spin column-type total RNA purification kit (NucleoSpin RNA; Takara Bio, Otsu, Japan) according to the manufacturer’s instructions. The MGIEasy RNA Directional Library Prep Set (MGI Tech) was used to prepare a complementary DNA library for next-generation sequencing of the extracted RNA. RNA sequencing was performed using a DNBSEQ-G400 (MGI Tech).

The genome sequences of *K. phaffii* GS115 were used as reference sequences for read mapping using Geneious prime version 2020.0.3 (Tomy Digital Biology, Tokyo, Japan). Differentially expressed genes (DEGs) were identified by calculating differential expression log2 ratios and p-values using Geneious. DEGs in the mutant strain were screened using the composite criteria of a p-value < 0.01 and at least a 1.5-fold change in expression (which means log2 [fold change] is ≥ 0.59, or ≤ −0.59). RNA sequencing data were deposited in the DDBJ nucleotide sequence database under the accession number PRJDB19698.

## 3. Results

### 3.1 Selection and cultivation of mutant strains generated by UV mutagenesis of GS115/S8/Z3

UV mutagenesis has been used to enhance D-lactic acid production from methanol in yeast. Specifically, UV irradiation was applied to the D-lactic acid-producing yeast strain GS115/S8/Z3, resulting in the isolation of 1,080 colonies, which were designated as DLac-Mut1-1 to DLac-Mut1-1080. These mutant strains were cultured in 1.0 mL of YPM medium using a microplate. Culture supernatants were collected after 48 h of cultivation and D-lactic acid concentrations were measured.

The D-lactic acid concentrations of the parent strain GS115/S8/Z3 and the 1,080 mutant strains after 48 h of cultivation are presented in Fig. 1. The D-lactic acid concentration in the parental strain GS115/S8/Z3 was 2.19 g/L. Among the mutant strains, 431 strains showed improved D-lactic acid production. From these 431 strains, the top seven strains with the highest D-lactic acid concentrations were selected: DLac_Mut1_89 (4.28 g/L), DLac_Mut1_372 (4.15 g/L), DLac_Mut1_129 (4.06 g/L), DLac_Mut1_25 (4.00 g/L), DLac_Mut1_260 (3.99 g/L), DLac_Mut1_322 (3.91 g/L), and DLac_Mut1_1077 (3.87 g/L).

**Fig. 1.**
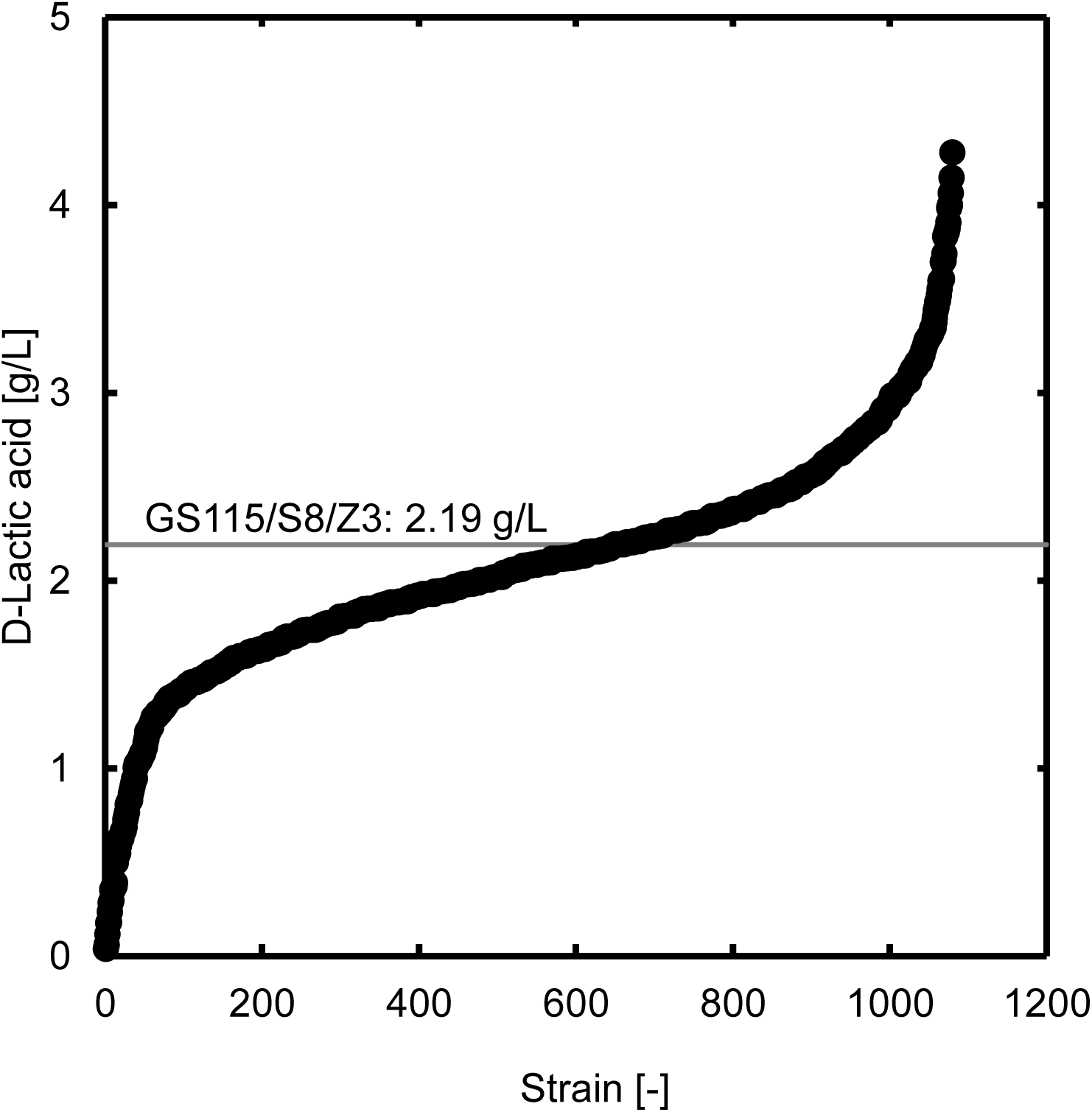
D-Lactic acid concentration after 48-h microplate cultivation of mutants obtained by UV mutagenesis of GS115/S8/Z3. Values are displayed in ascending order from left to right.

Fig. 2 shows the time-course changes in methanol concentration (Fig. 2(A)), optical density (OD_600_) (Fig. 2(B)), and D-lactic acid concentration (Fig.2(C)) during the flask cultivation of the seven selected mutant strains and the parent strain GS115/S8/Z3. The maximum D-lactic acid production by the seven selected mutant strains and the parent strain is shown in Fig. 2(D). As shown in Fig. 2(A), DLac-Mut1-89 and DLac-Mut1-129 exhausted methanol at 168 h, whereas the other strains depleted it by 144 h. Fig. 2(B) shows that the OD_600_ values for all strains increased significantly up to 48 h of cultivation, reaching ∼30, after which the change in cell density became minimal. There were no substantial differences in cell growth between the parent strain and the seven mutant strains. As shown in Figs 4 (C) and (D), compared with the parental strain, the four mutant strains (DLac_Mut1_25, DLac_Mut1_129, DLac_Mut1_322, and DLac_Mut1_1077) showed statistically significant improvements in D-lactic acid production. Notably, DLac_Mut1_25 achieved the highest D-lactic acid concentration of 4.45 g/L after 120 h of cultivation, representing a 1.25-fold increase compared to the parent strain GS115/S8/Z3.

**Fig. 2.**
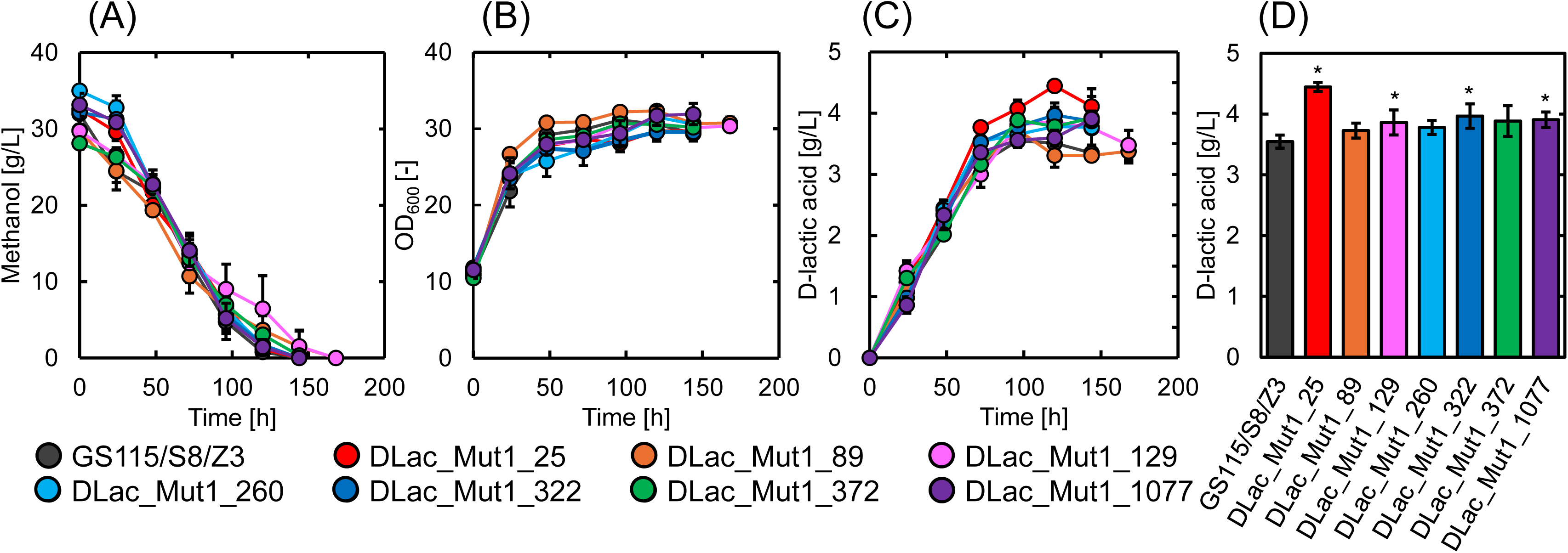
Time-course changes in (A) methanol concentration, (B) OD_600_, (C) D-lactic acid concentration, and (D) maximum D-lactic acid production in GS115/S8/Z3 and seven mutants. Data are presented as the average of three independent experiments. Error bars represent mean ± standard deviation. Statistical significance of the different D-lactic acid levels in comparison with the parent strain GS115/S8/Z3 was evaluated using Student’s t-test (*, P < 0.05).

### 3.2 Selection and cultivation of mutant strains generated by UV mutagenesis of DLac-Mut1-25

To further improve D-lactic acid production, we applied the same mutagenesis procedure to the mutant strain DLac_Mut1_25, which demonstrated enhanced D-lactic acid production after a single mutagenesis (Fig. 2). Overall, 1,080 mutant strains were isolated and designated DLac-Mut2-1 to DLac-Mut2-1080. These mutant strains were cultured in YPM medium (1.0 mL of YPM medium for 48 h using a microplate), and D-lactic acid concentrations were measured.

D-lactic acid concentrations in the 1,080 mutant strains after 48 h of cultivation are shown in Fig. 3. The D-lactic acid concentration in DLac-Mut1-25 during microplate cultivation was 4.00 g/L. Among the mutant strains, 10 strains exhibited higher D-lactic acid concentrations. The top eight strains were selected: DLac-Mut2-602 (4.50 g/L), DLac_Mut2_741 (4.43 g/L), DLac_Mut2_566 (4.32 g/L), DLac_Mut2_697 (4.28 g/L), DLac_Mut2_946 (4.24 g/L), DLac_Mut2_857 (4.20 g/L), DLac_Mut2_819 (4.20 g/L), and DLac_Mut2_221 (4.09 g/L).

**Fig. 3.**
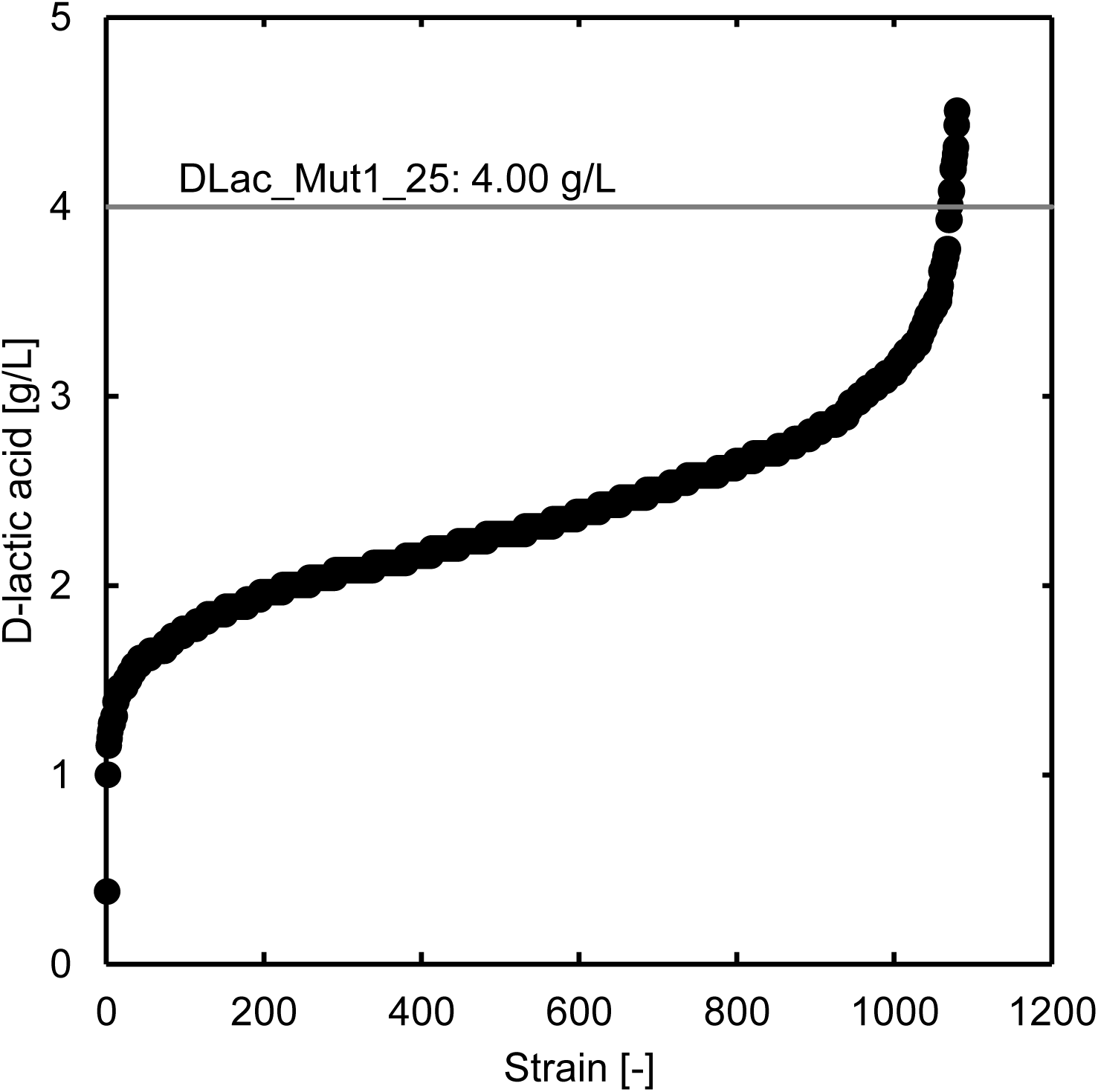
D-Lactic acid concentration after 48-h microplate cultivation of mutants obtained by UV mutagenesis of DLac_Mut1_25. Values are displayed in ascending order from left to right.

**Fig. 4.**
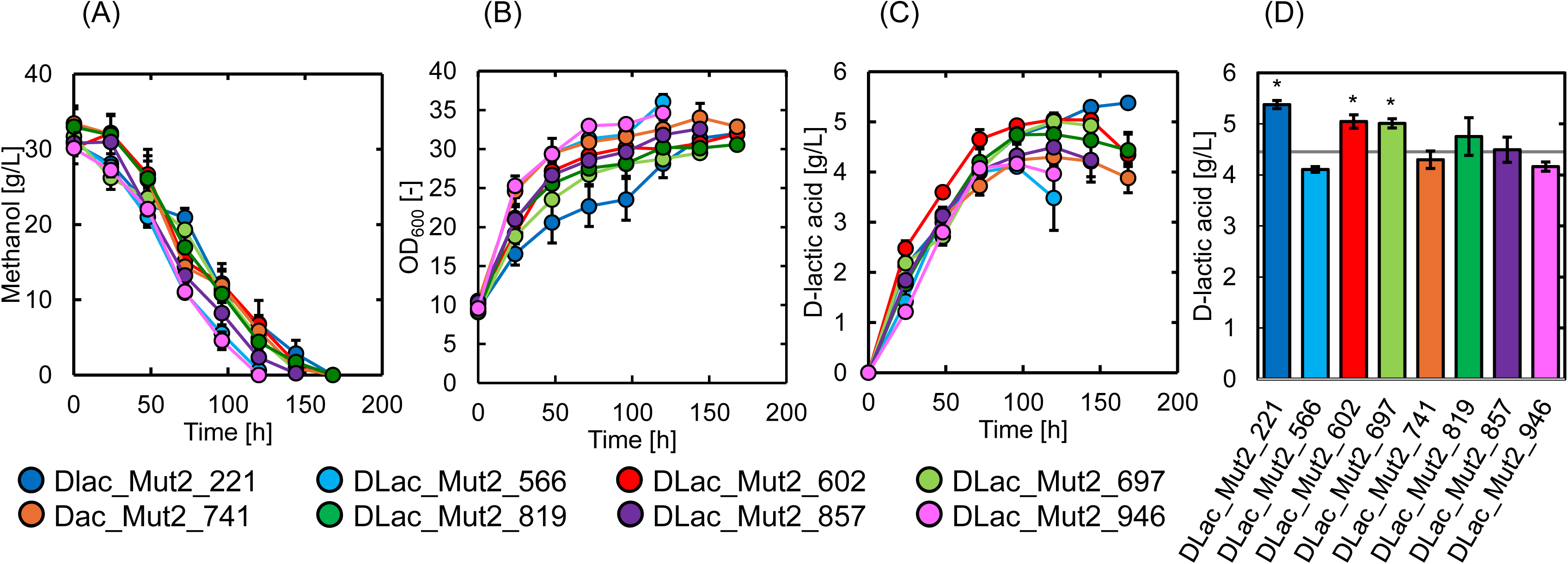
Time-course changes in (A) methanol concentration, (B) OD_600_, (C) D-lactic acid concentration, and (D) maximum D-lactic acid production of eight mutants. Gray line in (D) indicates D-lactic acid production in DLac_Mut1_25 (4.45 g/L). Data are presented as the average of three independent experiments. Error bars represent mean ± standard deviation. Statistical significance of the different D-lactic acid levels in comparison with the DLac_Mut1_25 was evaluated using Student’s t-test (*, P < 0.05).

Fig 4 shows the time-course changes in methanol concentration (Fig.4(A)), optical density (OD_600_) (Fig. 4(B)), and D-lactic acid concentration (Fig.4(C)) during flask cultivation of these eight mutant strains. Maximum D-lactic acid production by the eight mutant strains is shown in Fig. 4(D). As shown in Fig. 4(A), methanol depletion occurred at different time points: DLac_Mut2_566 and DLac_Mut2_946 at 120 h; DLac_Mut2_857 and DLac_Mut2_697 at 144 h; and DLac_Mut2_602, DLac_Mut2_741, DLac_Mut2_221, and DLac_Mut2_819 at 168 h. As shown in Fig. 4(B), DLac_Mut2_566 and DLac_Mut2_946 exhibited higher OD_600_ values than the other strains with significantly elevated cell densities after 120 h of cultivation.

DLac_Mut2_221 showed lower OD_600_ values until 96 h, but subsequently reached levels comparable to those of the other strains. As shown in Figs (C) and (D), compared to DLac-Mut1-25, three strains (DLac_Mut2_221, DLac_Mut2_602, and DLac_Mut2_697) showed statistically significant improvements in D-lactic acid production. DLac_Mut2_221 produced the highest D-lactic acid concentration of 5.38 g/L after 168 h of cultivation. This represents a 1.21-fold increase compared to DLac-Mut1-25 (4.45 g/L) and a 1.52-fold increase compared to the parent strain GS115_S8/Z3 (3.55 g/L).

### 3.3 Transcriptome analysis of the D-lactic acid production-improved mutant strain DLac_Mut2_221

Transcriptome analysis was performed to elucidate the cause of increased D-lactic acid production in the mutant strain DLac_Mut2_221. The -log10 (p-value), which indicates statistical significance, and log2 (fold change), which indicates changes in gene expression levels, were calculated for each gene and are shown as volcano plots (Fig.5). Genes with a high -log10 (p-value) were considered reliable, and genes with a high log2 (fold change) showed higher transcription levels in the mutant strain DLac_Mut2_221 than in the parent strain GS115/S8/Z3.

**Fig. 5.**
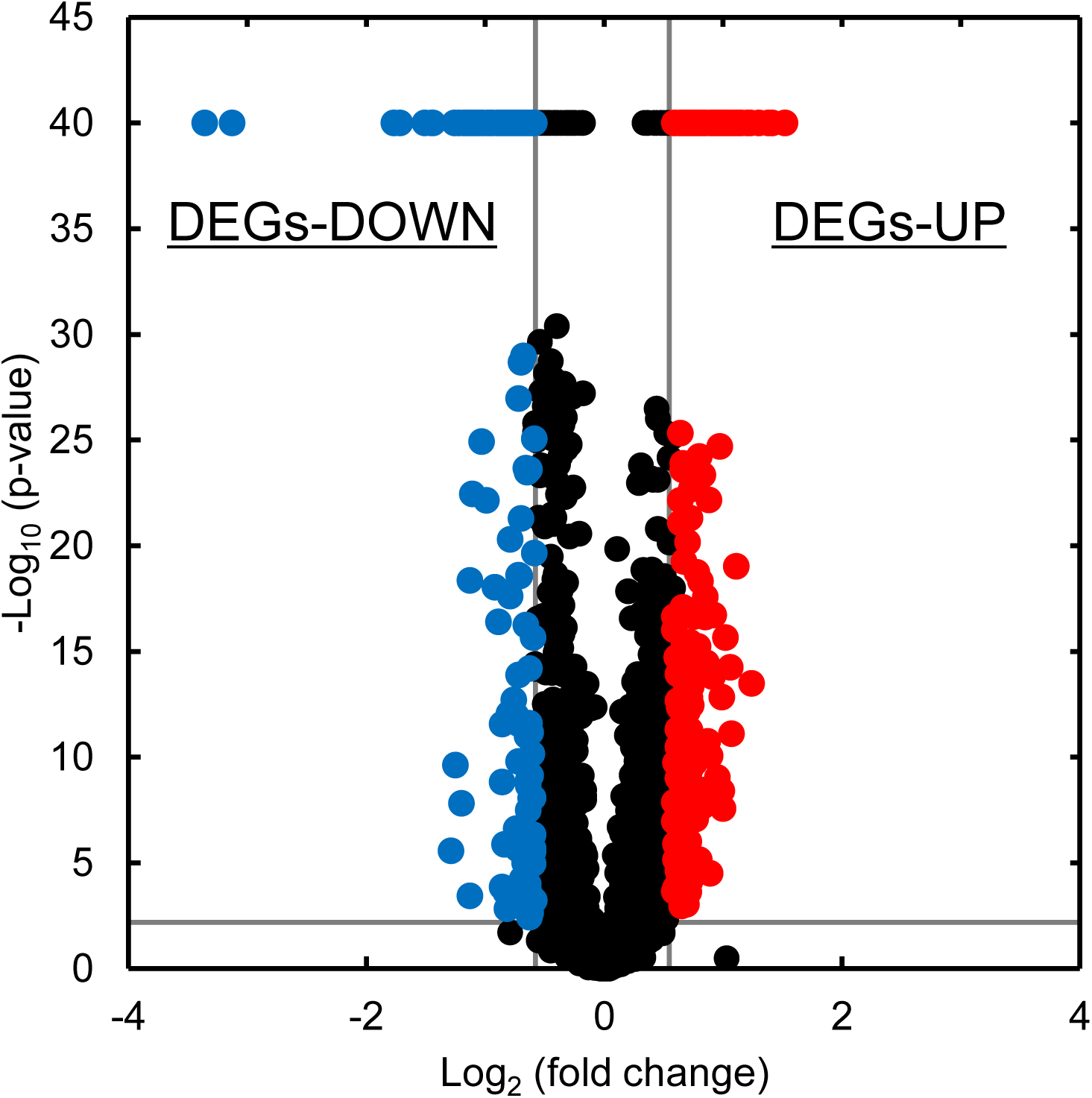
Volcano plot of genes for mutant Dlac_Mut2_221. Vertical lines represent ± 1.5-fold change (log2 fold change = ± 0.59). Horizontal line indicates a p-value of 0.01 (-log10 p-value =2.0). Red and blue plots represent DEG-UP and DEG-DOWN, respectively, in yeast. Black plots represent non-DEGs (i.e., genes with no change in transcription level).

The transcriptome analysis revealed 167 DEGs in DLac_Mut2_221, comprising 104 upregulated and 63 downregulated genes (Supplementary Tables S1 and S2). Among the identified DEGs, the top 20 upregulated and downregulated genes with known functions are summarized in Tables 1 and 2, respectively. Genes with increased expression in DLac_Mut2_221 included NAD(+)-dependent formate dehydrogenase, which catalyzes the dissimilation of formate to CO_2_, and malate synthase, which is involved in the glyoxylate cycle. Conversely, the downregulated genes included those involved in cell wall formation, such as *O-* glycosylated proteins required for cell wall stability.

**Table 1.** Top 20 genes of DLac_Mut2_221 with fold change upregulated via mutagenesis.

| Gene products | Locus tag | Fold change | -Log10 (p-value) |
| --- | --- | --- | --- |
| Glycerol kinase | PAS_chr4_0783 | 2.87 | 40.00 |
| Putative channel-like protein | PAS_chr4_0784 | 2.66 | 40.00 |
| Glycerol kinase | PAS_chr4_0783 | 2.55 | 40.00 |
| Putative channel-like protein | PAS_chr4_0784 | 2.35 | 40.00 |
| Thiazole synthase | PAS_chr3_0648 | 2.35 | 40.00 |
| Malate synthase | PAS_chr4_0191 | 2.30 | 40.00 |
| Protein involved in synthesis of the thiamine precursor hydroxy methylpyrimidine (HMP) | PAS_chr4_0065 | 2.17 | 40.00 |
| Putative transmembrane protein involved in export of ammonia, a starvation signal | PAS_chr1-1_0158 | 2.17 | 40.00 |
| Nuclear protein related to mammalian high mobility group (HMG) proteins | PAS_chr4_0796 | 2.14 | 40.00 |
| Protein containing an N-terminal epsin-like domain | PAS_chr4_0763 | 2.06 | 40.00 |
| Acetate--CoA ligase | PAS_chr2-1_0767 | 2.01 | 40.00 |
| Ubiquitin | PAS_chr4_0762 | 1.92 | 40.00 |
| Cytochrome c, isoform 1 | PAS_chr4_0018 | 1.85 | 40.00 |
| Essential nucleolar protein involved in the early steps of 35S rRNA processing | PAS_chr4_0274 | 1.85 | 10.06 |
| Hexokinase-2 | PAS_chr1-4_0561 | 1.80 | 40.00 |
| Thioredoxin peroxidase | PAS_chr2-2_0220 | 1.80 | 40.00 |
| H subunit of the mitochondrial glycine decarboxylase complex | PAS_FragB_0009 | 1.79 | 40.00 |
| Plasma membrane localized protein that protects membranes from desiccation | PAS_chr4_0627 | 1.79 | 40.00 |
| Mitochondrial 54S ribosomal protein YmL39 | PAS_chr3_1135 | 1.78 | 40.00 |
| NAD(+)-dependent formate dehydrogenase | PAS_chr3_0932 | 1.78 | 40.00 |

**Table 2.** Top 20 genes of DLac_Mut2_221 with fold change down regulated via mutagenesis.

| Gene products | Locus tag | Fold change | -Log10 (p-value) |
| --- | --- | --- | --- |
| O-glycosylated protein required for cell wall stability | PAS_chr4_0305 | 0.35 | 40.00 |
| C-8 sterol isomerase | PAS_chr4_0198 | 0.37 | 40.00 |
| Putative chitin transglycosidase, cell wall protein | PAS_chr4_0559 | 0.43 | 40.00 |
| Putative chitin transglycosidase, cell wall protein | PAS_chr4_0820 | 0.46 | 40.00 |
| Mitochondrial outer membrane and cell wall localized SUN family member | PAS_chr4_0559 | 0.47 | 40.00 |
| Pho85 cyclin of the Pcl1,2-like subfamily | PAS_chr2-1_0092 | 0.49 | 18.15 |
| Ammonium permease | PAS_chr1-4_0394 | 0.53 | 11.35 |
| O-glycosylated protein required for cell wall stability | PAS_chr4_0305 | 0.53 | 40.00 |
| Aromatic aminotransferase II | PAS_chr4_0147 | 0.54 | 40.00 |
| 60S ribosomal protein L22 | PAS_chr4_0041 | 0.58 | 40.00 |
| Major exo-1,3-beta-glucanase of the cell wall | PAS_chr2-1_0454 | 0.58 | 15.28 |
| Mitochondrial intermembrane space protein | PAS_chr4_0561 | 0.59 | 8.20 |
| Vacuolar cation channel | PAS_chr4_0220 | 0.59 | 11.46 |
| Protoporphyrinogen oxidase | PAS_chr4_0055 | 0.59 | 12.71 |
| 40S ribosomal protein S6 | PAS_chr4_0094 | 0.61 | 40.00 |
| ADP/ATP carrier protein | PAS_chr4_0210 | 0.61 | 40.00 |
| Cytoplasmic RNA-binding protein, contains an RNArecognition motif (RRM) | PAS_chr2-1_0414 | 0.61 | 7.88 |
| Bud neck-localized, SH3 domain-containing protein required for cytokinesis | PAS_chr2-2_0259 | 0.64 | 9.04 |
| Polyamine transport protein specific for spermine | PAS_chr1-3_0215 | 0.64 | 11.92 |
| Vacuolar carboxypeptidase yscS | PAS_chr4_0686 | 0.64 | 40.00 |

## 4. Discussion

Through two-step UV mutagenesis, we obtained the mutant strain DLac_Mut2_221, which produced 5.38 g/L of D-lactic acid (1.52-fold higher than that of the parent strain) after 168 h of culture. (Fig. 4(D)). In our previous study, we successfully produced 5.18 g/L of D-lactic acid from methanol in *K. phaffii* by selecting the *DLDH* gene, optimizing the expression promoter, and integrating multiple copies into the rDNA locus (Inoue et al., 2024). Sae-Tang et al. also reported that expression of the *DLDH* gene from *Leuconostoc pseudomesenteroides* and the flocculation gene from *Saccharomyces cerevisiae* in *K. phaffii* improved D-lactic acid production from glucose (Sae-Tang et al., 2023). Wu *et al*. demonstrated the production of 4.2 g/L L-lactic acid by introducing mutations in the *LLDH* gene from *Lactobacillus plantarum*, inhibiting lactic acid consumption pathways, and localizing the enzyme to the mitochondria (Wu et al., 2025). To the best of our knowledge, the DLac_Mut2_221 mutant strain was generated by simple UV mutagenesis and achieved the highest lactic acid production concentrations, including D- and L-lactic acid, from methanol as the sole carbon source. This represents an advancement in D-lactic acid production from methanol using *K. phaffii* and demonstrates the effectiveness of UV mutagenesis as a strategy for strain improvement in *K. phaffii*.

UV mutagenesis has the advantage that it is easy to obtain a variety of mutant strains and that changes in gene expression can be easily investigated by genomic and transcriptomic analyses of mutant strains. These benefits have led to research on UV mutagenesis in various microorganisms. Weng *et al* reported that S-adenosyl-L-methionine production in *S. cerevisiae* is associated with enhanced ATP synthesis through the reinforcement of the TCA cycle and gluconeogenesis/glycolysis pathways (Weng et al., 2022). Similarly, Kamba et al. demonstrated that genes related to glyceraldehyde 3-phosphate and dihydroxyacetone phosphate synthesis contributed to improved triacylglycerol production in yeast *Lipomyces starkeyi* (Kamba et al., 2024). If a phenotype can be enhanced by random mutagenesis, such as UV mutagenesis, a detailed analysis of the mutant strains can be used to identify previously unnoticed genes that affect the target phenotype. Furthermore, the identified genes can be manipulated by overexpression or deletion to further enhance the desired phenotype. Even in modern metabolic engineering, where the latest technologies such as genome editing are frequently used, random mutagenesis is still considered useful.

Among the genes upregulated in DLac_Mut2_221, formate dehydrogenase (*FDH*) showed a notable 1.78-fold increase in expression (Table 1 and Fig. 6). Additionally, slight increases in the S-formylglutathione hydrolase (*FGH*) (1.17-fold) and formaldehyde dehydrogenase (*FLD*) (1.42-fold) levels were observed (Fig. 6). These enzymes are part of the formaldehyde dissimilation pathway (Duman et al., 2020; Yu et al., 2022). In *K. phaffii*, Jia *et al*. previously discussed formaldehyde toxicity and successfully improved monellin production by suppressing formaldehyde accumulation (Jia et al., 2022). Similarly, in another methanol-utilizing microorganism, *Pichia methanolica*, studies have revealed that formaldehyde was more toxic than methanol during metabolism (Wakayama et al., 2016). Increased expression of *FDH*, *FGH*, and *FLD* likely enhances the formaldehyde dissimilation pathway. Notably, the expression of *AOX*, which converts methanol to formaldehyde, decreased slightly (0.74-fold) (Fig. 6). These gene expression changes suggest a potential mechanism by which methanol conversion to formaldehyde is suppressed, while the formaldehyde dissimilation pathway is simultaneously strengthened, thereby avoiding formaldehyde accumulation. This mechanism likely contributes to the improved D-lactic acid production from methanol by mitigating formaldehyde-associated toxicity.

**Fig. 6.**
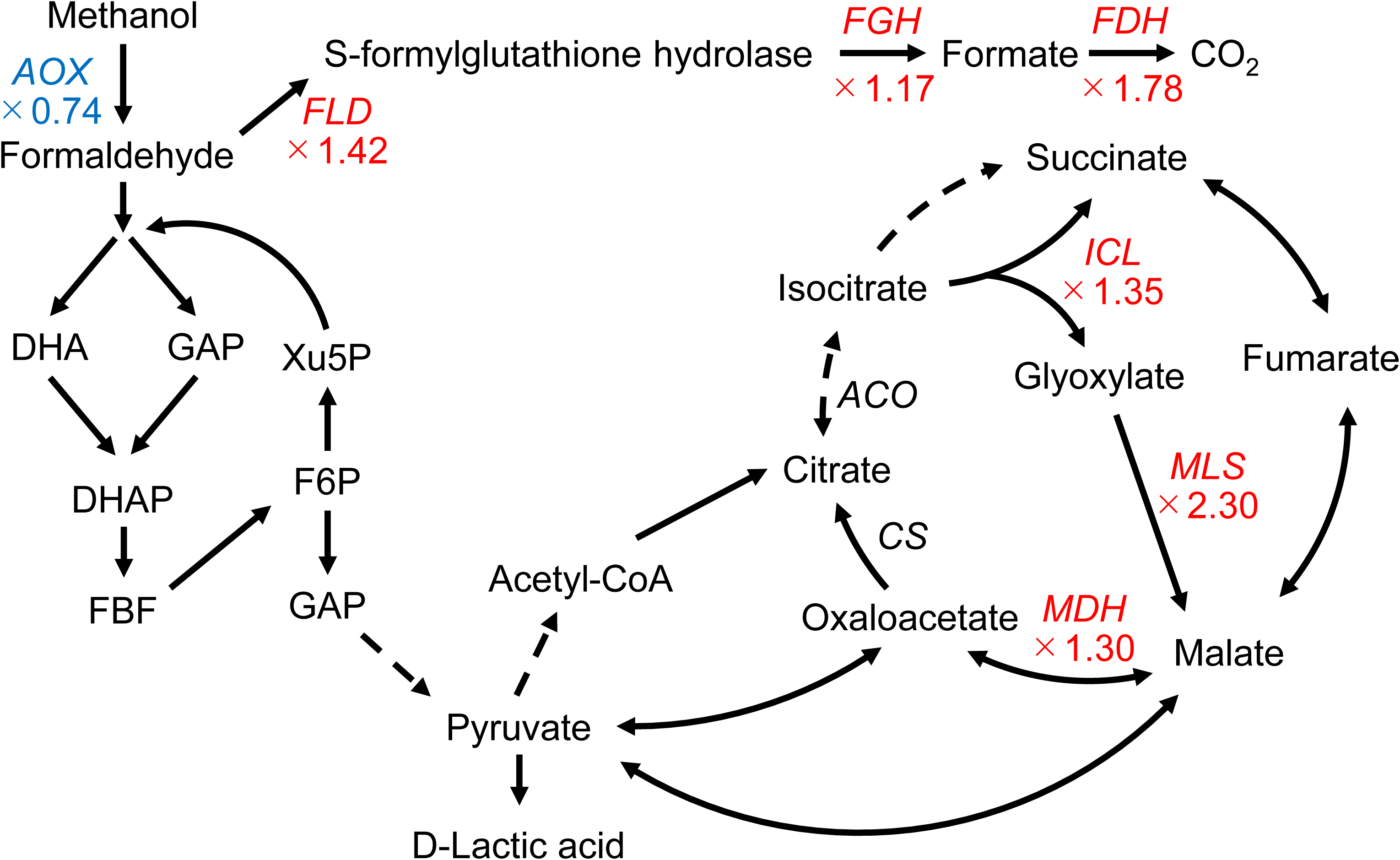
Schematic of methanol utilization pathway in *K. phaffii*. *ACO*: Aconitase, *AOX*: Alcohol oxidase; *CS*: Citrate synthase; DHA: Dihydroxyacetone, DHAP: Dihydroxyacetone Phosphate, F6P: Fructose-6-phosphate, FBP: Fructose-1,6-Bisphosphate, *FDH*: Formate dehydrogenase; *FGH*: S-formylglutathione hydrolase; *FLD*: Formaldehyde dehydrogenase; GAP: Glyceraldehyde 3-phosphate; *ICL*: Isocitrate lyase; *MDH*: Malate dehydrogenase; *MLS*: Malate synthase; Xu5P: Xylulose 5-phosphate. Solid lines represent a single reaction, whereas dashed lines represent multiple reactions. Enzyme genes marked in red represent upregulated genes, whereas those marked in blue represent downregulated genes.

In the DLac_Mut2_221 strain, malate synthase (*MLS*), an enzyme that converts glyoxylate to malate, showed a 2.30-fold increase in expression (Table 1 and Fig. 6). This enzyme is involved in the glyoxylate pathway. Additionally, other glyoxylate pathway-related genes showed increased expression. Isocitrate lyase (*ICL*), which produces malate and succinate from isocitrate, showed a 1.35-fold increase (Fig. 6), and malate dehydrogenase (*MDH*), which catalyzes the conversion of malate to oxaloacetate, showed a 1.30-fold increase (Fig. 6). The glyoxylate pathway is comprised of five key enzymes: *ICL*, *MLS*, *MDH*, citrate synthase (*CS*), and aconitase (*ACO*) (Kunze et al., 2002). The primary enzymes, *ICL* and *MLS*, convert isocitrate into succinate and malate, respectively. Although structurally similar to the TCA cycle, the glyoxylate pathway is distinct because it does not release CO_2_, thereby enabling carbon conservation (Kunze et al., 2006). In *K. phaffii*, the glyoxylate pathway is minimally utilized when glucose is used as the carbon source. However, when methanol is used as the carbon source, the glyoxylate pathway is activated, facilitating the synthesis of C4 compounds (Rußmayer et al., 2015). Based on these observations, DLac_Mut2_221 demonstrated an increased expression of glyoxylate pathway-related genes, suggesting a more robust activation of the glyoxylate pathway. This enhanced activation likely prevents carbon loss through CO_2_ conversion in the TCA cycle, thereby potentially improving the conversion of carbon to D-lactic acid.

The gene with the most significantly decreased expression in the DLac_Mut2_221 strain was the *O*-glycosylated protein required for cell wall stability (PAS_0305), which decreased 0.35-fold (Table 2). Guo et al. reported that deletion of PAS_0305 increased cell wall permeability, leading to improved protein expression and CMCase activity (Gao et al., 2023). Furthermore, flux balance analysis revealed that carbon loss is reduced in the PAS_0305-deficient strain (Gao et al., 2023). Therefore, in DLac_Mut2_221, the decrease in PAS_0305 gene expression likely contributed to improved D-lactic acid production through reduced carbon loss.

## 5 Conclusions

In this study, UV mutagenesis of the metabolically engineered D-lactic acid-producing *K. phaffii* GS115/S8/Z3 yielded the mutant strain DLac_Mut2_221, which was capable of efficiently producing D-lactic acid from methanol. The mutant strain was confirmed to produce 5.38 g/L of D-lactic acid from 30 g/L methanol after 168 h of cultivation. To the best of our knowledge, this is the highest concentration of lactic acid, including D- and L-lactic acid, produced from methanol as the sole carbon source. Transcriptome analysis of the mutant strain revealed that three important mechanisms were presumably involved in the increased D-lactic acid production from methanol in the mutant strain: (1) avoiding excessive formaldehyde accumulation, (2) activating the glyoxylate pathway, and (3) reducing the expression of the *O*-glycosylated protein required for cell wall stability. To the best of our knowledge, there are no previous reports linking activation of the glyoxylate pathway or reduced expression of the *O*-glycosylated protein required for cell wall stability to compound production from methanol in *K. phaffii*. This study suggests that these genes may contribute to improving the production of valuable compounds from methanol. Currently, there is an increasing amount of research on the production of useful compounds from methanol and various approaches have been proposed to improve the production of target compounds from methanol. The knowledge gained from this study, combined with that gained from previous research on improving compound productivity, will further improve the productivity of the target compounds from methanol.

---

**Table S1.**
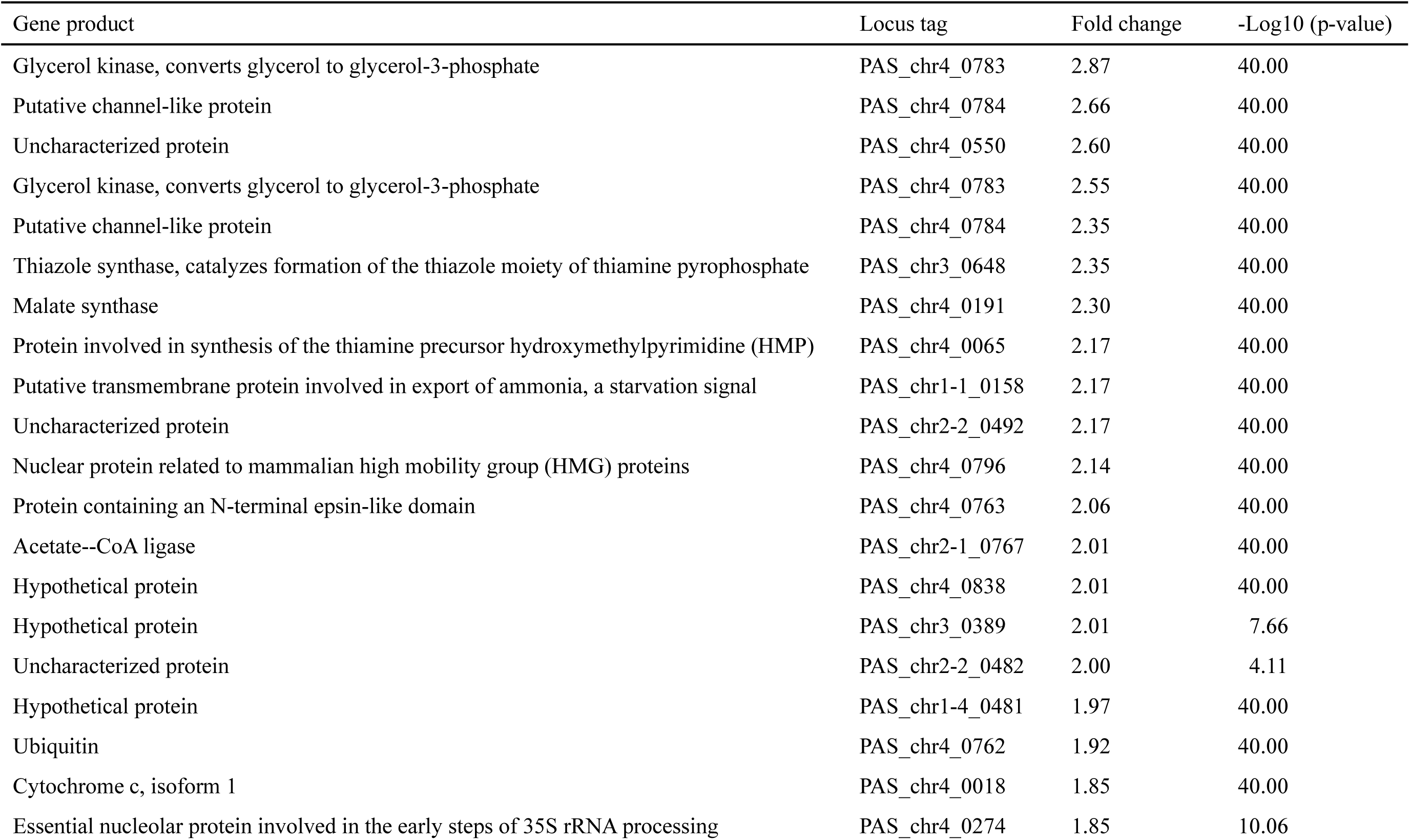
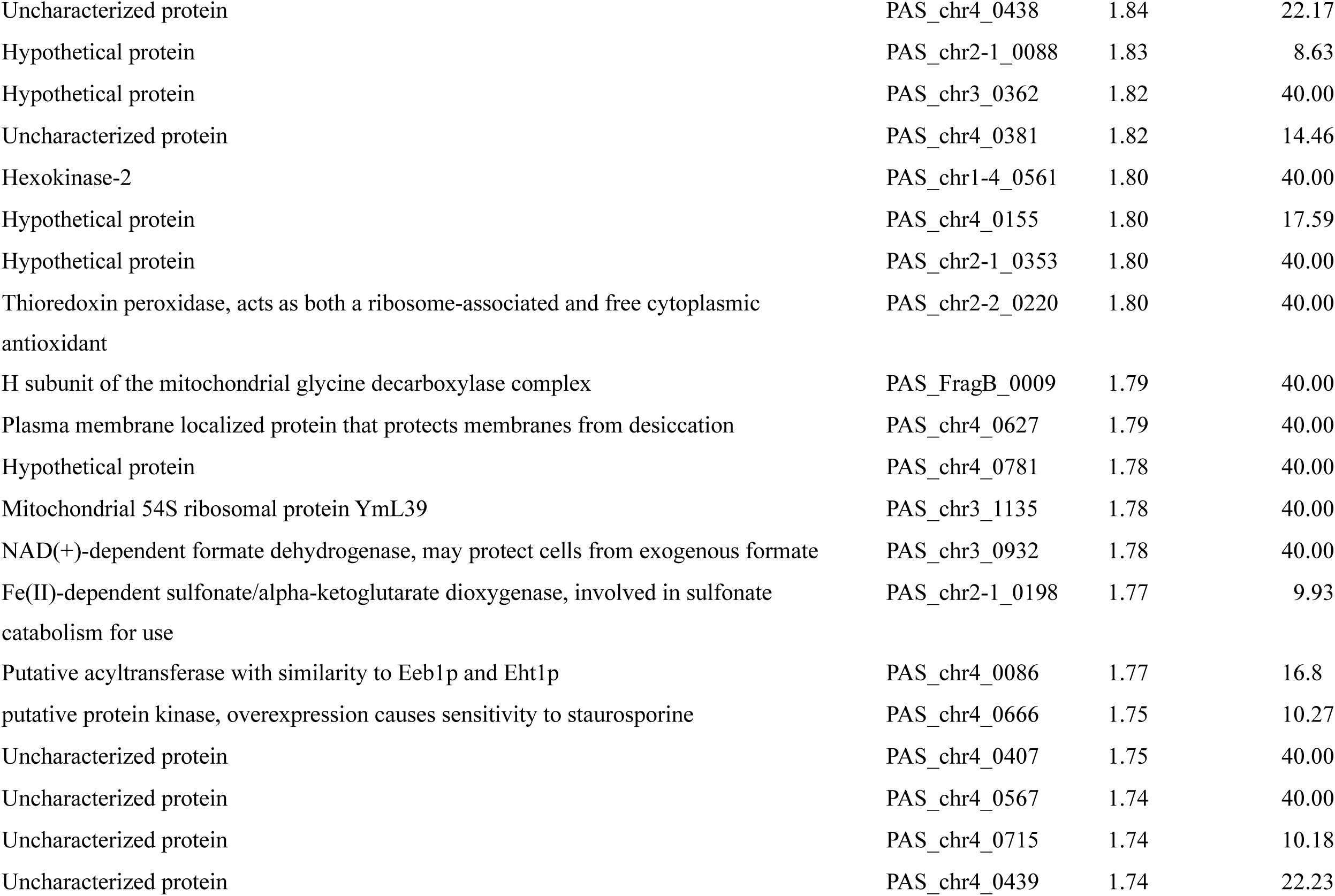
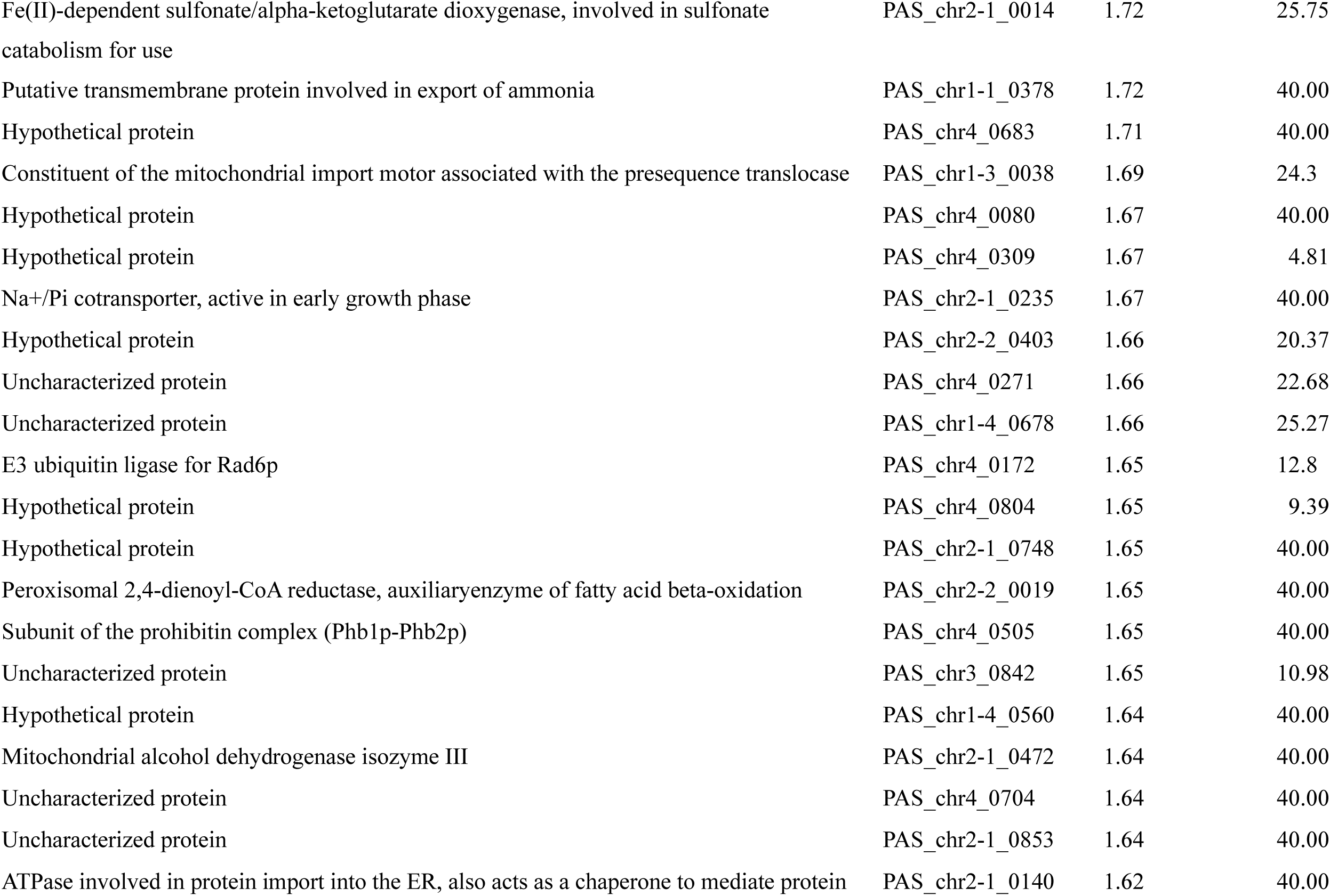
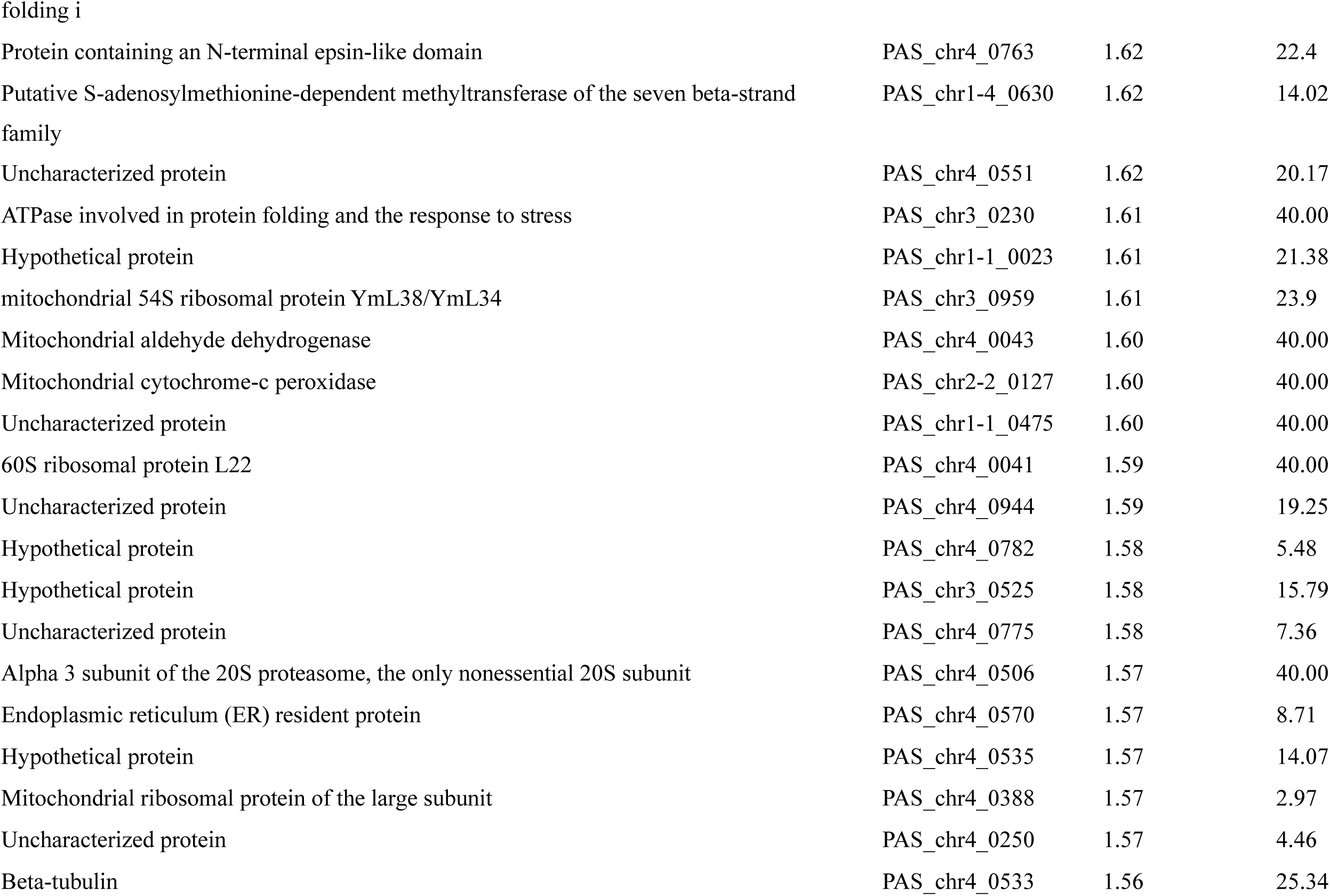
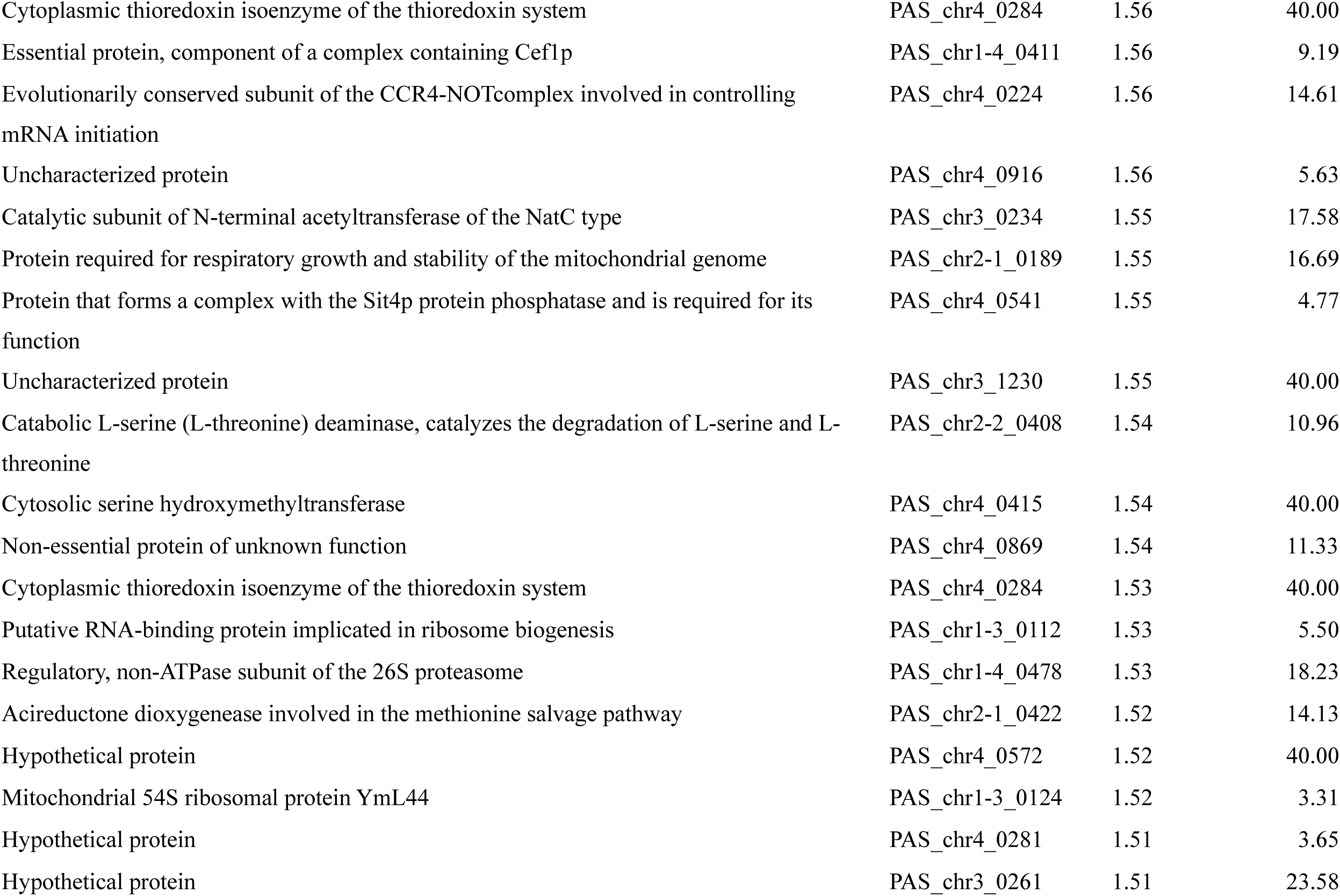
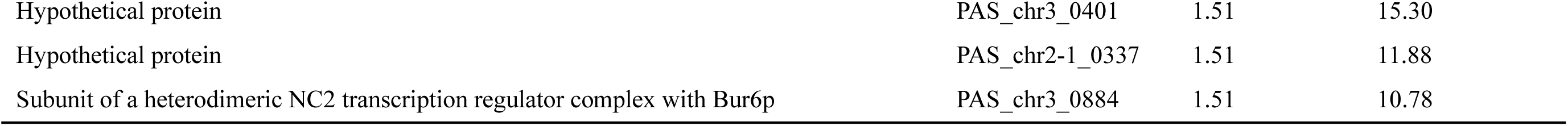
Upregulated genes in the mutant strain DLac_Mut2_221.

**Table S2.** Downregulated genes in the mutant strain DLac_Mut2_221.

| Gene product | Locus tag | Fold change | -Log10 (p-value) |
| --- | --- | --- | --- |
| Hypothetical protein | PAS_chr4_0151 | 0.11 | 40.00 |
| Uncharacterized protein | PAS_chr4_0820 | 0.29 | 40.00 |
| Hypothetical protein | PAS_chr2-1_0002 | 0.33 | 40.00 |
| O-glycosylated protein required for cell wall stability | PAS_chr4_0305 | 0.35 | 40.00 |
| C-8 sterol isomerase | PAS_chr4_0198 | 0.37 | 40.00 |
| Putative chitin transglycosidase, cell wall protein | PAS_chr4_0559 | 0.43 | 40.00 |
| Uncharacterized protein | PAS_chr4_0820 | 0.43 | 40.00 |
| Hypothetical protein | PAS_chr4_0461 | 0.46 | 18.37 |
| Hypothetical protein | PAS_chr4_0553 | 0.46 | 40.00 |
| Putative chitin transglycosidase, cell wall protein | PAS_chr4_0559 | 0.46 | 40.00 |
| Mitochondrial outer membrane and cell wall localized SUN family member | PAS_chr4_0046 | 0.47 | 40.00 |
| Uncharacterized protein | PAS_chr4_0914 | 0.47 | 40.00 |
| Pho85 cyclin of the Pcl1,2-like subfamily, involved in entry into the mitotic cell cycle and regulat | PAS_chr2-1_0092 | 0.49 | 18.15 |
| Uncharacterized protein | PAS_chr4_0926 | 0.51 | 40.00 |
| Uncharacterized protein | PAS_chr2-2_0489 | 0.51 | 40.00 |
| Ammonium permease involved in regulation of pseudohyphal growth | PAS_chr1-4_0394 | 0.53 | 11.35 |
| Hypothetical protein | PAS_chr4_0461 | 0.53 | 12.21 |
| O-glycosylated protein required for cell wall stability | PAS_chr4_0305 | 0.53 | 40.00 |
| Uncharacterized protein | PAS_chr1-1_0479 | 0.54 | 40.00 |
| Aromatic aminotransferase II | PAS_chr4_0147 | 0.54 | 40.00 |
| Hypothetical protein | PAS_chr1-4_0327 | 0.55 | 40.00 |
| Uncharacterized protein | PAS_chr1-1_0482 | 0.55 | 40.00 |
| Hypothetical protein | PAS_chr4_0089 | 0.55 | 11.56 |
| Major exo-1,3-beta-glucanase of the cell wall, involved in cell wall beta-glucan assembly | PAS_chr2-1_0454 | 0.58 | 15.28 |
| Hypothetical protein | PAS_chr4_0114 | 0.58 | 17.63 |
| 60S ribosomal protein L22 | PAS_chr4_0041 | 0.58 | 40.00 |
| Hypothetical protein | PAS_chr1-4_0554 | 0.58 | 3.87 |
| Hypothetical protein | PAS_chr1-1_0234 | 0.58 | 21.63 |
| Uncharacterized protein | PAS_chr2-2_0488 | 0.58 | 40.00 |
| Mitochondrial intermembrane space protein, forms a complex with TIM8p | PAS_chr4_0561 | 0.59 | 8.2 |
| uncharacterized protein | PAS_chr1-3_0290 | 0.59 | 8.68 |
| Vacuolar cation channel | PAS_chr4_0220 | 0.59 | 11.46 |
| Protoporphyrinogen oxidase | PAS_chr4_0055 | 0.59 | 12.71 |
| Uncharacterized protein | PAS_chr4_0460 | 0.60 | 5.47 |
| Uncharacterized protein | PAS_chr4_0779 | 0.61 | 4.54 |
| Uncharacterized protein | PAS_FragB_0059 | 0.61 | 5.64 |
| Hypothetical protein | PAS_chr4_0335 | 0.61 | 11.58 |
| 40S ribosomal protein S6 | PAS_chr4_0094 | 0.61 | 40.00 |
| Hypothetical protein | PAS_chr1-4_0583 | 0.61 | 2.26 |
| Cytoplasmic RNA-binding protein, contains an RNA recognition motif (RRM) | PAS_chr2-1_0414 | 0.61 | 7.88 |
| ADP/ATP carrier protein | PAS_chr4_0210 | 0.61 | 40.00 |
| Hypothetical protein | PAS_chr4_0628 | 0.62 | 21.30 |
| Hypothetical protein | PAS_chr4_0152 | 0.62 | 40.00 |
| Uncharacterized protein | PAS_chr1-3_0303 | 0.62 | 8.74 |
| Hypothetical protein | PAS_chr4_0590 | 0.63 | 16.26 |
| uncharacterized protein | PAS_chr4_0972 | 0.63 | 40.00 |
| Hypothetical protein | PAS_chr1-4_0240 | 0.64 | 3.85 |
| Bud neck-localized, SH3 domain-containing protein required for cytokinesis | PAS_chr2-2_0259 | 0.64 | 9.04 |
| Hypothetical protein | PAS_chr4_0553 | 0.64 | 10.83 |
| Uncharacterized protein | PAS_chr3_0076 | 0.64 | 40.00 |
| Hypothetical protein | PAS_chr2-2_0117 | 0.64 | 10.98 |
| Polyamine transport protein specific for spermine | PAS_chr1-3_0215 | 0.64 | 11.92 |
| Hypothetical protein | PAS_chr4_0883 | 0.64 | 40.00 |
| Vacuolar carboxypeptidase yscS | PAS_chr4_0686 | 0.64 | 40.00 |
| Uncharacterized protein | PAS_chr4_0919 | 0.65 | 11.60 |
| Putative dihydrokaempferol 4-reductase | PAS_chr4_0336 | 0.65 | 14.19 |
| G1/S-specific cyclin | PAS_FragB_0046 | 0.66 | 4.52 |
| Meiosis-specific protein of unknown function, required for spore wall formation during sporulation | PAS_chr4_0560 | 0.66 | 5.54 |
| L-homoserine-O-acetyltransferase | PAS_chr2-1_0383 | 0.66 | 6.96 |
| Uncharacterized protein | PAS_chr1-3_0299 | 0.66 | 7.27 |
| Uncharacterized protein | PAS_chr4_0915 | 0.66 | 8.09 |
| Uncharacterized protein | PAS_chr3_1239 | 0.66 | 2.02 |
| Uncharacterized protein | PAS_chr2-1_0887 | 0.66 | 40.00 |
| Protein of the SUN family (Sim1p, Uth1p, Nca3p, Sun4p) that may participate in DNA replication | PAS_chr2-2_0064 | 0.67 | 27.65 |

